# Cyrius: accurate *CYP2D6* genotyping using whole genome sequencing data

**DOI:** 10.1101/2020.05.05.077966

**Authors:** Xiao Chen, Fei Shen, Nina Gonzaludo, Alka Malhotra, Cande Rogert, Ryan J Taft, David R Bentley, Michael A Eberle

## Abstract

Responsible for the metabolism of 25% of clinically used drugs, *CYP2D6* is a critical component of personalized medicine initiatives. Genotyping *CYP2D6* is challenging due to sequence similarity with its pseudogene paralog *CYP2D7* and a high number and variety of common structural variants (SVs). Here we describe a novel bioinformatics method, Cyrius, that accurately genotypes *CYP2D6* using whole-genome sequencing (WGS) data. We show that Cyrius has superior performance (96.5% concordance with truth genotypes) compared to existing methods (84-86.8%). After implementing the improvements identified from the comparison against the truth data, Cyrius’s accuracy has since been improved to 99.3%. Using Cyrius, we built a haplotype frequency database from 2504 ethnically diverse samples and estimate that SV-containing star alleles are more frequent than previously reported. Cyrius will be an important tool to incorporate pharmacogenomics in WGS-based precision medicine initiatives.

## Introduction

There is significant variation in the response of individuals to a large number of clinically prescribed drugs. A strong contributing factor to this variability in drug metabolism is the genetic composition of the drug-metabolizing enzymes, and thus genotyping pharmacogenes is important for personalized medicine^1^. Cytochrome P450 2D6 (*CYP2D6*) encodes one of the most important drug-metabolizing enzymes and is responsible for the metabolism of about 25% of clinically used drugs^2^. The *CYP2D6* gene is highly polymorphic, with 131 star alleles defined by the Pharmacogene Variation (PharmVar) Consortium^3^. Star alleles^4^ are *CYP2D6* haplotypes defined by a combination of small variants (SNVs and indels) and structural variants (SVs), and correspond to different levels of *CYP2D6* enzymatic activity, i.e. poor, intermediate, normal, or ultrarapid metabolizer^5–7^.

Genotyping *CYP2D6* is challenged by common deletions and duplications of *CYP2D6* and fusions (commonly referred to as hybrids) between *CYP2D6* and its pseudogene paralog, *CYP2D7*^4,8,9^, which shares 94% sequence similarity, including a few near-identical regions^8,10^. The interrogation of SVs improves the accuracy of *CYP2D6* phenotype prediction^11^. Traditionally, *CYP2D6* genotyping is done in low or medium throughput with array-based platforms, such as the PharmacoScan, or polymerase chain reaction (PCR) based methods such as TaqMan assays, ddPCR and long-range PCR. These assays differ in the number of star alleles (variants) they interrogate, leading to variability in genotyping results across assays^8,12,13^. To detect SVs, these conventional platforms need to be complemented with CNV assays that may also be limited to detection of just a subset of the known CNVs^4,9^.

With recent advances in next-generation sequencing (NGS), it is now possible to profile the entire genome at high-throughput and in a clinically-relevant timeframe. Driven by these advances, many countries are undertaking large scale population sequencing projects^14–16^ wherein pharmacogenomics testing will greatly increase the clinical utility of these efforts. There exists a few bioinformatics tools for genotyping *CYP2D6* (Cypiripi^17^, Astrolabe (formerly Constellation)^18^, Aldy^19^ and Stargazer^20,21^) that can be applied to targeted (PGRNseq^22^) and/or whole genome sequencing (WGS) data. Among these, Cypiripi and Astrolabe (both developed more than four years ago) were not designed to detect complex SVs and have been shown to have lower performance than the more recently developed methods^19,23,24^. The two most recent *CYP2D6* callers, Aldy and Stargazer, work by detecting SVs based on sequencing coverage and calling star alleles based on the observed small variants and SVs. They rely on accurate read alignments, which may not be possible at many positions throughout the gene as the sequence is highly similar or even indistinguishable with *CYP2D7*. Relying on the initial read alignments may lead to ambiguous read coverage patterns or false positive/negative small variant calls. Another limitation of both Aldy and Stargazer is that, currently, neither method supports the hg38 genome build so studies using the latest genome build (hg38) will require a re-alignment to hg19/GRCh37 to use these tools.

Here we describe Cyrius, a novel WGS-specific *CYP2D6* genotyping tool that overcomes the challenges with the homology between *CYP2D6* and *CYP2D7* and works for sequence data aligned to both hg38 and hg19. The availability of a panel of reference samples by the CDC Genetic Testing Reference Material Program (GeT-RM)^12,25^, where the consensus genotypes of major pharmacogenes are derived using multiple genotyping platforms, has enabled assessment of genotyping accuracy for newly developed methods. Furthermore, the recent availability of high-quality long reads can provide a complete picture of *CYP2D6* for improved validation of complicated variants and haplotypes^25,26^. We demonstrate superior genotyping accuracy compared to other methods in 138 GeT-RM reference samples and 8 samples with PacBio HiFi data, covering 40 known star alleles. We applied this method to WGS data on 2504 unrelated samples from the 1000 Genomes Project^27^ and report on the distribution of star alleles across five ethnic populations. This analysis expands the current understanding of the genetic diversity of *CYP2D6*, particularly on complex star alleles with SVs.

## Methods

### Samples

We included 138 GeT-RM reference samples in our truthset^12,25^. WGS was performed for 96 samples with TruSeq DNA PCR-free sample preparation and 2×150bp reads sequenced on Illumina HiSeq X instruments. Genome build GRCh37 was used for read alignment with Isaac v04.16.09.24^28^. The WGS data for the remaining 42 samples were downloaded as part of the 1000 Genomes Project (see below).

For population analysis, trio concordance tests and truthset comparison, we downloaded WGS BAM files from the 1000 Genomes Project (1kGP) (See Availability of Data and Materials). These BAM files were generated by sequencing 2×150bp reads on Illumina NovaSeq 6000 instruments from PCR-free libraries sequenced to an averaged depth of at least 30X and aligned to the human reference, hs38DH, using BWA-MEM v0.7.15.

PacBio sequencing data for 8 samples (Table S1) were downloaded from 1kGP and the Genome in a Bottle (GIAB) Consortium.

### *CYP2D6* genotyping method used by Cyrius

Read alignment accuracy is reduced in *CYP2D6* because of the homology with *CYP2D7* (Figure 1) and this can make variant calling challenging and error prone. Cyrius uses a novel approach to overcome this challenge and a detailed workflow is described below and illustrated in Figure 2 using NA12878 (*\*3/*68*+*\*4*) as an example.

**Figure 1.**
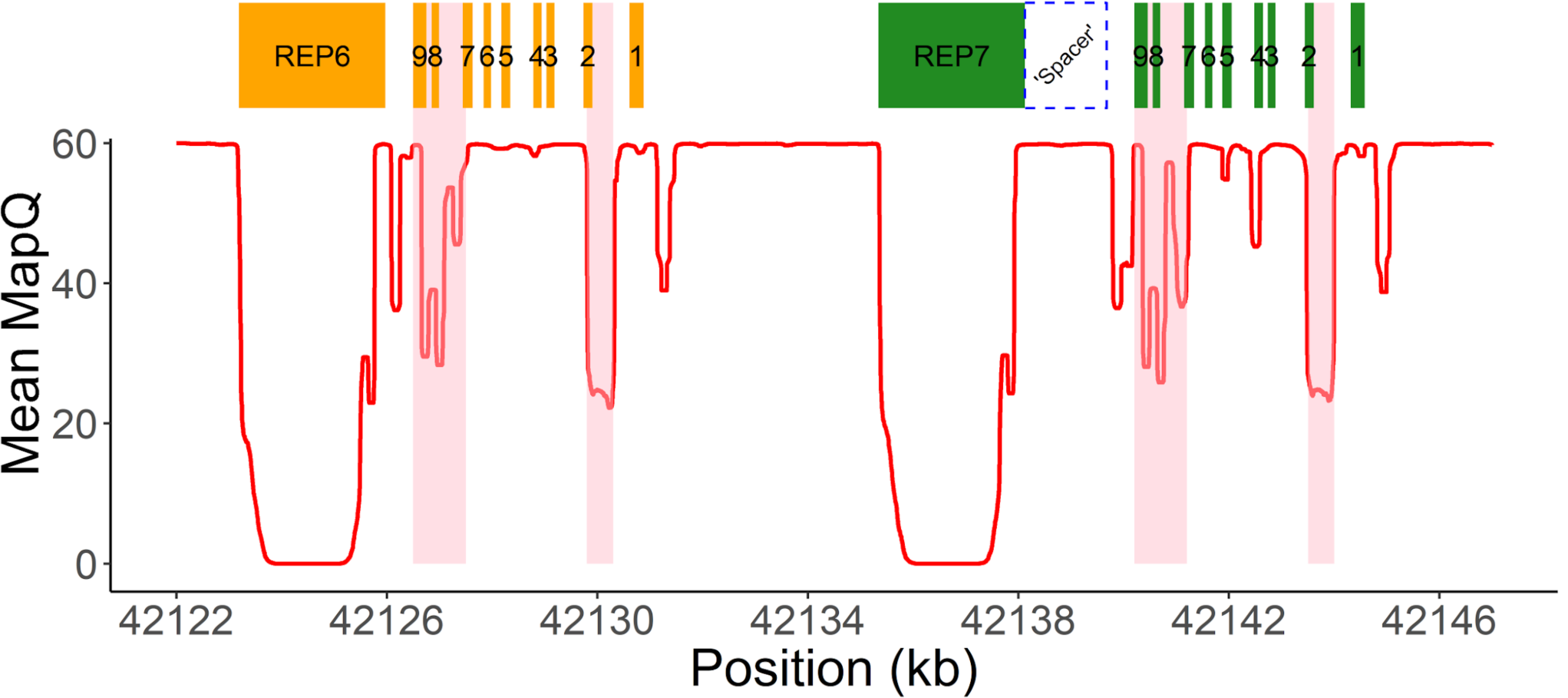
WGS data quality in *CYP2D6/CYP2D7* region. Mean mapping quality (red line) averaged across 1kGP samples plotted for each position in the *CYP2D6/CYP2D7* region (hg38). A median filter is applied in a 200bp window. The 9 exons of *CYP2D6/CYP2D7* are shown as orange (*CYP2D6*) and green (*CYP2D7*) boxes. Two 2.8kb repeat regions downstream of *CYP2D6* (REP6) and *CYP2D7* (REP7) are near-identical and essentially unalignable. The purple dashed line box denotes the unique spacer region between *CYP2D7* and REP7. Two major homology regions within the genes are shaded in pink and highlight areas of low mapping accuracy.

**Figure 2.**
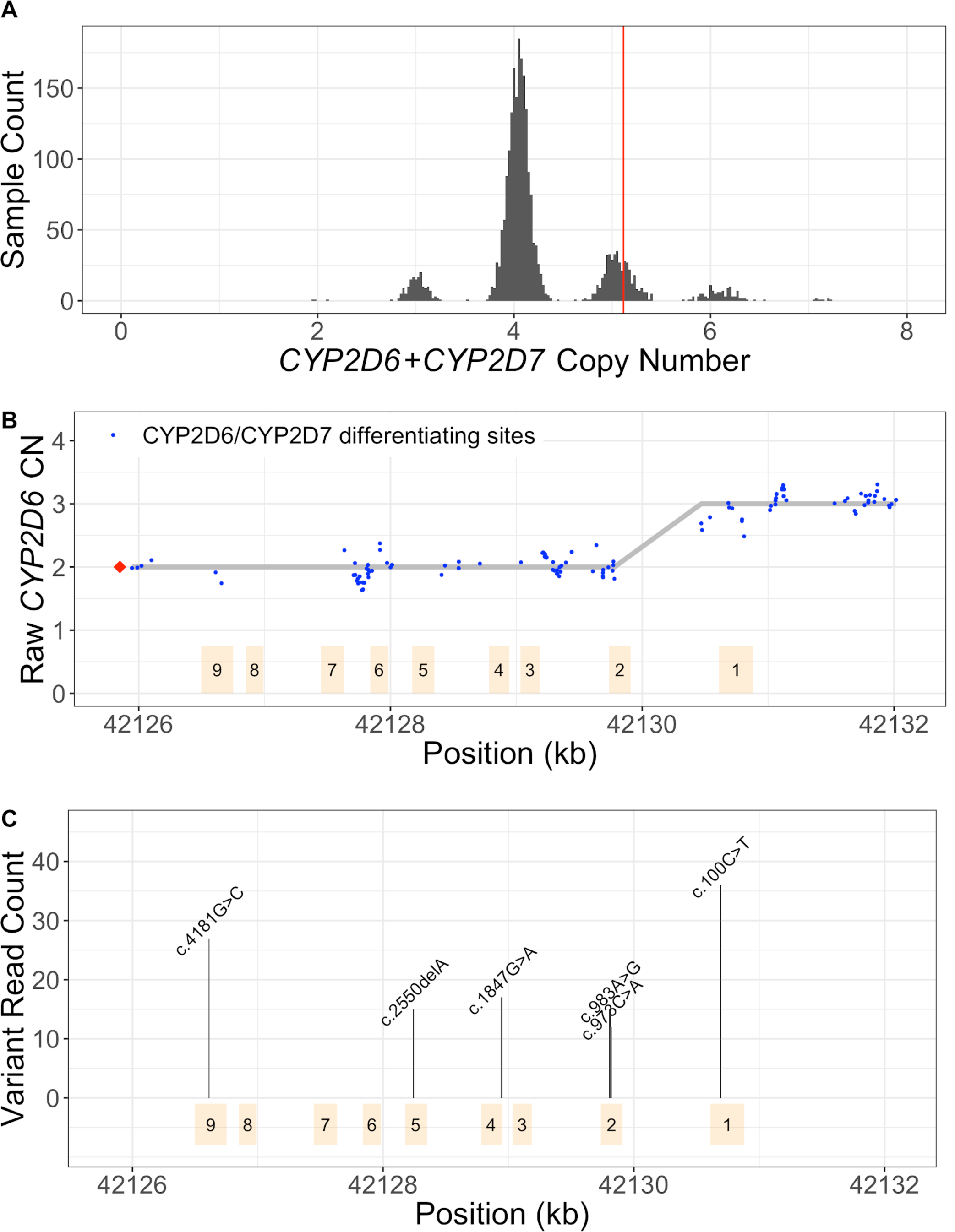
Cyrius workflow, using NA12878 (*\*3/*68*+*\*4*) as an example. **A**. CN(*CYP2D6*+*CYP2D7*) is derived by counting and modelling all reads that align to either *CYP2D6* or *CYP2D7*. The histogram shows the distribution of normalized *CYP2D6*+*CYP2D7* depth in 1kGP samples, showing peaks at CN2, 3, 4, 5, 6 and 7. The red vertical line represents the value for NA12878, corresponding to CN5 that indicates an additional copy (could be *CYP2D6* or hybrid). **B**. SVs are called by examining the CNs of *CYP2D6*/*CYP2D7* differentiating bases. Exons are denoted by yellow boxes. Blue dots denote raw *CYP2D6* CNs, calculated as CN(*CYP2D6*+*CYP2D7*) multiplied by the ratio of *CYP2D6* supporting reads out of *CYP2D6* and *CYP2D7* supporting reads. The red diamond denotes the CN of genes that are *CYP2D6*-derived at the 3’ end (can be complete *CYP2D6* or *CYP2D7-CYP2D6* hybrid), calculated as CN(*CYP2D6*+*CYP2D7*) minus CN(spacer). The *CYP2D6* CN is called at each *CYP2D6/CYP2D7* differentiating site and a change in *CYP2D6* CN within the gene indicates the presence of a hybrid. In NA12878, the *CYP2D6* CN changes from 2 to 3 between Exon 2 and Exon 1, indicating a *CYP2D6*-*CYP2D7* hybrid (*\*68*). **C**. Supporting read counts of the star-allele defining protein-changing small variants are used to call the CN of each variant. The y axis shows the read counts for all queried small variant positions. Six variants are called in NA12878, one of which, c.100C>T, is called as two copies (one copy belongs to *\*4* and the other belongs to *\*68*). Finally, star alleles are called based on detected SVs and small variants.

First, Cyrius identifies the total copies of both *CYP2D6* and *CYP2D7* combined (i.e. CN(*CYP2D6*+*CYP2D7*)) following a similar method as previously described^27^. Read counts are calculated directly from the WGS aligned BAM file using all reads mapped to either *CYP2D6* or *CYP2D7*, including reads aligned with a mapping quality of zero because of the homology between *CYP2D6* and *CYP2D7*. The summed read count is normalized and corrected for GC content and CN(*CYP2D6*+*CYP2D7*) is called from a Gaussian mixture model built on the normalized depth values. While there exist ambiguous alignments between *CYP2D6* and *CYP2D7*, the sequencing coverage for both genes combined exhibits a clean signal (Figure 2A), allowing us to identify SVs that result in a gain or loss in CN(*CYP2D6*+*CYP2D7*). CN(*CYP2D6*+*CYP2D7*) is 4 in samples without SV; a CN(*CYP2D6*+*CYP2D7*) of 3 suggests a deletion of either *CYP2D6* or *CYP2D7*; a CN(*CYP2D6*+*CYP2D7*) of 5 suggests an extra copy, which could be *CYP2D6* duplication or a hybrid. The red vertical line in Figure 2A shows the results for NA12878 where we identified five copies of *CYP2D6* plus *CYP2D7*. We use the same approach to call the CN of the 1.6kb spacer region between the repeat REP7 and *CYP2D7* (Figure 1). The CN(spacer) indicates the summed CN of *CYP2D7* and *CYP2D6*-*CYP2D7* hybrids. Thus, subtracting CN(spacer) from CN(*CYP2D6*+*CYP2D7*) gives the summed CN of *CYP2D6* and *CYP2D7*-*CYP2D6* hybrids.

We next determine the number of complete *CYP2D6* genes as well as identify hybrid genes. To do this we identified 117 reliable bases that differ between *CYP2D6* and *CYP2D7* (Supplementary Information and Figure S1) and use these to identify the exact form of SVs that impact *CYP2D6*. Cyrius estimates the *CYP2D6* CN at each of the 117 *CYP2D6*/*CYP2D7* differentiating base positions. Based on CN(*CYP2D6*+*CYP2D7*), Cyrius calls the combination of *CYP2D6* CN and *CYP2D7* CN that produces the highest likelihood for the observed number of reads supporting *CYP2D6-* and *CYP2D7*-specific bases, as described previously^29^. Hybrids are identified when the CN of *CYP2D6* changes within the gene. For example, NA12878 shown in Figure 2B has two full copies of *CYP2D6* and one hybrid where Exon 1 comes from *CYP2D7* and Exons 2-9 come from *CYP2D6* (i.e. *\*68*).

Next Cyrius parses the read alignments to identify the protein-changing small variants that define star alleles and call their CNs (Figure 2C). These variants are divided into two classes: 1) variants that fall in *CYP2D6/CYP2D7* homology regions, i.e. the shaded low mapping quality regions in Figure 1, and 2) variants that occur in unique regions of *CYP2D6*. For the former, Cyrius looks for variant reads in *CYP2D6* and its corresponding site in *CYP2D7* to account for possible mis-alignments, i.e. a *CYP2D6* read that aligns to *CYP2D7*. For the latter, Cyrius only uses the reads aligned to *CYP2D6*. The total *CYP2D6* CN at the variant sites are taken into account during small variant calling so that a variant can be called at one copy, two copies or any CN less than or equal to the *CYP2D6* CN at that site.

Finally, Cyrius matches the SVs and small variants against star allele definitions (PharmVar, last accessed on 7/15/2020) and produces star allele calls in diplotypes that are consistent with the called variants (Supplemental Information).

### Validating against truth from GeT-RM and long reads

We confirmed that all the star alleles in our validation data are interrogated by Cyrius, Aldy and Stargazer. When comparing the calls made by Cyrius, Aldy and Stargazer against the truth genotypes, a genotype is considered a match as long as all star alleles in the truth genotype are present, even if the haplotype assignment is different. For example, several samples listed in GeT-RM as *\*1/*10*+*\*36*+*\*36* are called by Aldy as *\*1*+*\*36/*10*+*\*36* and we considered these to be correct.

When validating genotype calls against the PacBio data, PacBio reads that cover the entire *CYP2D6* gene (one single haplotype) were analyzed to identify small variants and the corresponding star allele. Reads carrying SVs were determined by aligning reads against a set of reference contigs that were constructed to represent known SVs (*\*5, *13, *36, *68* and duplications). Visualization in Figure 4 was done using the software tool sv-viz2^30^.

**Figure 3.**
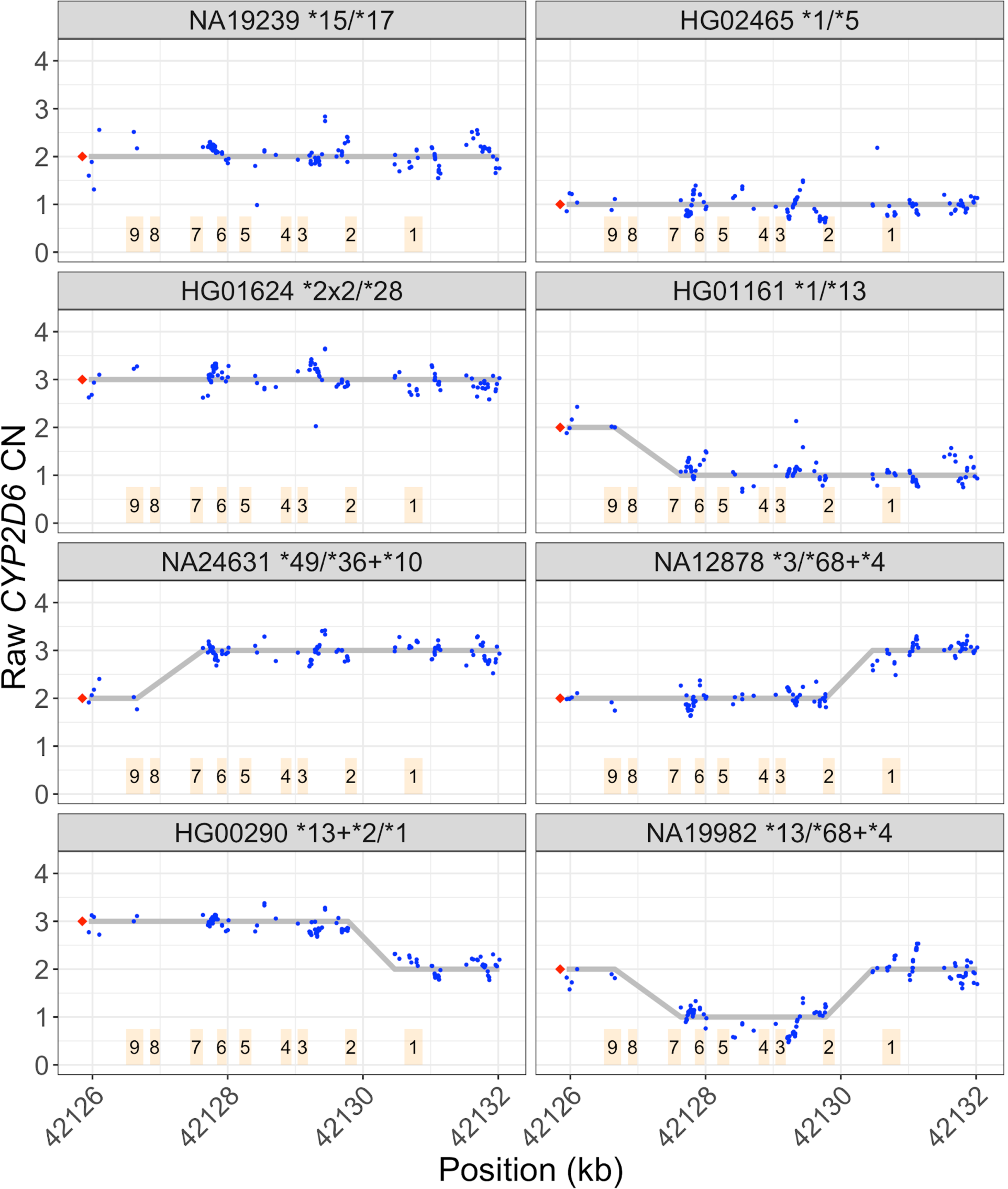
Depth patterns in samples with different types of SVs. Depth plots as described in Figure 2B. *CYP2D6* CN is called at each *CYP2D6/CYP2D7* differentiating site and a change in *CYP2D6* CN within the gene indicates the presence of a hybrid. The depth profiles for different SV patterns are shown in NA19239 (no SV), HG02465 (deletion, *\*5*), HG01624 (duplication), HG01161 (*CYP2D7-CYP2D6* hybrid, *\*13*), NA24631 (*CYP2D6-CYP2D7* hybrid, *\*36*), NA12878 (*CYP2D6-CYP2D7* hybrid, \**68*), HG00290 (tandem arrangement *\*13*+*\*2*), and NA19982 (two different SVs, *\*13* and *\*68*, one on each haplotype). The hybrids in NA24631 and NA12878 are confirmed with PacBio reads in Figure 4.

**Figure 4.**
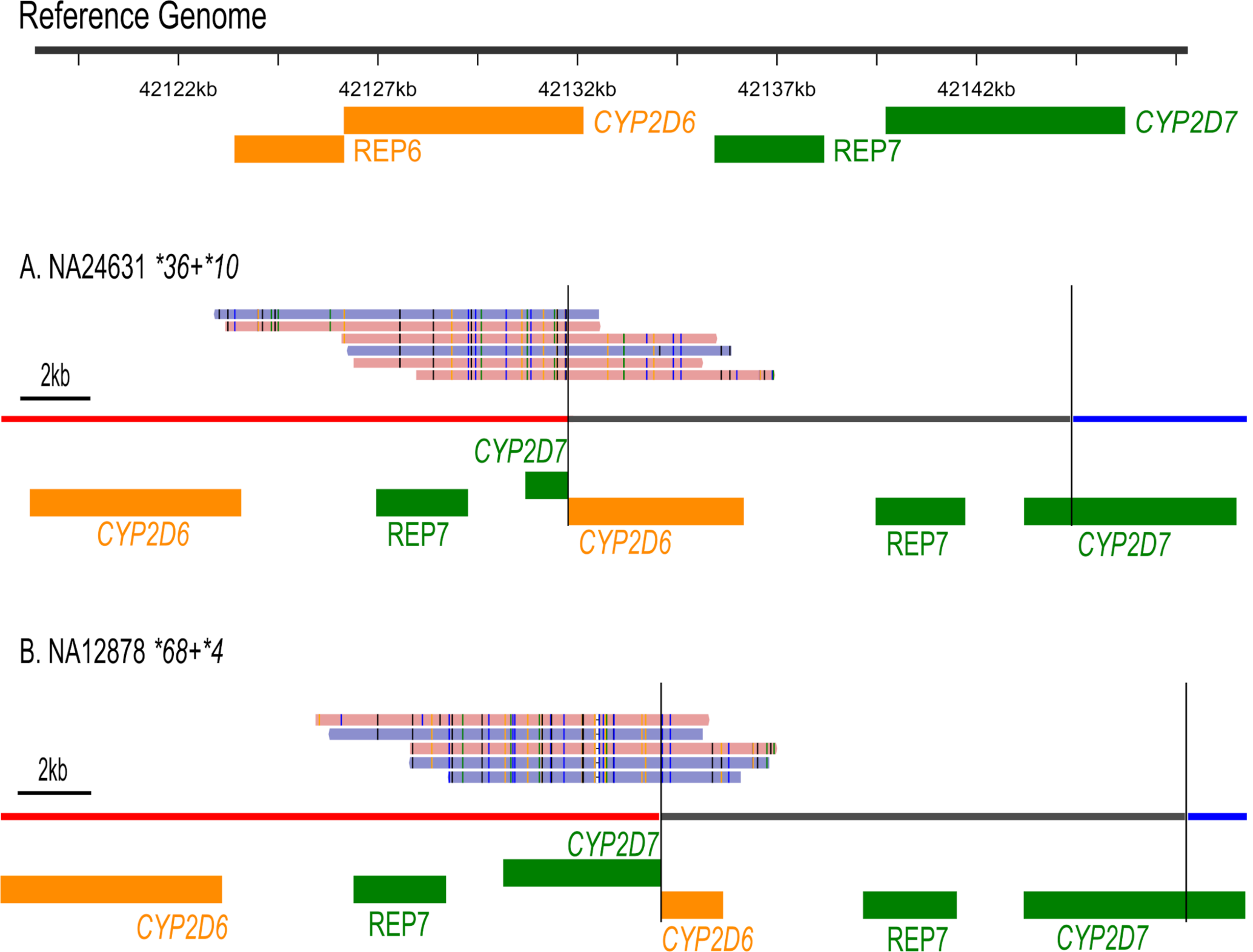
Structural variants validated by PacBio HiFi reads. PacBio reads supporting *CYP2D6*-*CYP2D7* hybrid *\*36* and *\*68*, confirming SVs called in NA24631 and NA12878 (third row, Figure 2). PacBio reads were realigned against modified sequence contigs representing the fusions and plotted using sv-viz2^22^. The black vertical lines mark the boundaries of the duplicated sequences, represented by the gray region (the original copy is in the red region). The red and blue regions represent flanking sequences.

### Running Aldy and Stargazer

Aldy v2.2.5 was run using the command “aldy genotype -p illumina -g *CYP2D6*”.

Stargazer v1.0.7 was run to genotype *CYP2D6* using VDR as the control gene, with GDF and VCF files as input.

The 1kGP GeT-RM samples were originally aligned against hg38. Though Cyrius is designed to work on either GRCh38 or GRCh37 (hg19), because Aldy and Stargazer currently only support GRCh37, for comparison between methods, these samples were realigned against GRCh37 using Isaac^28^.

## Results

### Validation and performance comparison

We compared the *CYP2D6* calls made by Cyrius, Aldy and Stargazer against 144 truthset samples, including 138 GeT-RM samples and 8 samples with truth generated using PacBio HiFi sequence reads (two samples overlap between GeT-RM and PacBio, Table S1). Samples with SVs show distinct depth signals that allow us to call SVs accurately (Figure 3). The long reads allow us to locate and visualize breakpoints of the common SVs in the region (Figure 4) and thus serve as a valuable resource for studying complex star alleles and confirm the phasing of the variants for the star alleles.

Comparing against the GeT-RM samples, we found three samples where the calls of all three software methods agree with each other but disagree with the GeT-RM consensus (Table S1). First, for NA18519, the WGS-based genotype is *\*106/*29* with reads carrying the variant defining *\*106* (Figure S2). This genotype is also confirmed by other studies^23,24^. The GeT-RM consensus is *\*1/*29*, because none of the GeT-RM assays interrogate *\*106* and the sample was not sequenced. The remaining two samples, NA23874 and NA24008, have the *\*68 CYP2D6-CYP2D7* hybrid that is not represented in the GeT-RM consensus. For these, the depth profiles show a CN gain in Exon 1 (Figure S3A) and PacBio long reads confirm the presence of *\*68* hybrid (Figure S3B/C). In GeT-RM testing, these two samples only underwent limited CNV testing (no TaqMan CNV result is available for Exon 1, the *CYP2D6* part of the hybrid). Therefore, based on this additional evidence, the GeT-RM truth genotypes for these two samples should be updated to include *\*68*. For the accuracy calculations below, we consider these three samples to be correctly genotyped by the WGS-based methods.

Cyrius initially made 5 discordant calls in the 144 truth samples, showing a concordance of 96.5% (Table 1). We were subsequently able to identify the causes and improve Cyrius to correctly call 4 of these 5 samples (Supplementary Information), reaching a ‘trained’ concordance of 99.3% (143 out of 144 samples). In contrast, both of the other *CYP2D6* callers had concordance less than 90%. Aldy had a concordance of 86.8% and, in particular, overcalled several hybrids such as *\*61, *63, *78* and *\*83* (called in 7 out of 19 discordant samples, Table S1), even in samples without SVs. Stargazer had a concordance of 84% and is most prone to errors when SVs are present. The concordance in samples with SVs is 75.9%, and 13 out of the 23 discordant calls are in samples with SVs (Table 1).

**Table 1.** Summary of benchmarking results against truth in 144 samples.

| Caller | Total concordant | Concordance | Deletion<br>N=15 | Duplication<br>N=13 | Hybrid<br>N=26 | No SV<br>N=90 | Concordance,<br>samples with<br>SV | Concordance,<br>samples<br>without SV |
| --- | --- | --- | --- | --- | --- | --- | --- | --- |
| Cyrius | 139* | 96.5% | 14 | 12 | 25 | 88 | 94.4% | 97.8% |
| Aldy | 125 | 86.8% | 13 | 11 | 23 | 78 | 87.0% | 86.7% |
| Stargazer | 121 | 84.0% | 14 | 10 | 17 | 80 | 75.9% | 88.9% |

\*Cyrius has since been improved and can correctly call 143 (99.3%) out of 144 of these samples.

Together, the validation samples used in this study confirmed our *CYP2D6* calling accuracy in 47 distinct haplotypes (<u>Table 2</u>), including 40 star alleles as well as several SV structures, such as duplications and tandem arrangements including *\*13*+*\*2, *68*+*\*4, *36*+*\*10 and *36*+*\*36*+*\*10*. Of these, *\*49* is not found in GeT-RM but present in a sample with PacBio data. These 40 star alleles represent 30.5% of the 131 star alleles in PharmVar and 51.7% (31 out of 60) of the star alleles with known function.

**Table 2.** Haplotypes validated in this study and their frequencies in 1kGP.

| Haplotype | Pan-ethnic<br>(N=2504) | European<br>(N=503) | Admixed-American<br>(N=347) | East-Asian<br>(N=504) | African<br>(N=661) | South-Asian<br>(N=489) | Validated in this<br>study | In GeT-RM<br>full set | Function |
| --- | --- | --- | --- | --- | --- | --- | --- | --- | --- |
| *1 | 33.35 | 35.88 | 45.97 | 26.09 | 25.87 | 39.37 | x | x | Normal |
| *2 | 14.76 | 15.9 | 18.59 | 7.74 | 12.71 | 20.86 | x | x | Normal |
| *3 | 0.54 | 1.79 | 0.58 | 0 | 0.23 | 0.2 | x | x | None |
| *4 | 5.93 | 11.83 | 9.22 | 0.2 | 2.34 | 8.28 | x | x | None |
| *5 | 3.49 | 2.39 | 2.02 | 3.47 | 5.82 | 2.56 | x | x | None |
| *6 | 0.5 | 2.09 | 0.29 | 0 | 0.08 | 0.1 | x | x | None |
| *7 | 0.18 | 0 | 0 | 0 | 0 | 0.92 | x | x | None |
| *9 | 0.68 | 2.39 | 1.3 | 0 | 0.08 | 0 | x | x | Decreased |
| *10 | 5.25 | 1.39 | 1.44 | 14.98 | 3.86 | 3.78 | x | x | Decreased |
| *11 | 0.02 | 0 | 0 | 0 | 0.08 | 0 | x | x | None |
| *13 | 0.1 | 0.2 | 0.14 | 0 | 0.08 | 0.1 | x | x | None |
| *14 | 0.18 | 0 | 0 | 0.89 | 0 | 0 | x | x | Decreased |
| *15 | 0.06 | 0 | 0 | 0 | 0.23 | 0 | x | x | None |
| *17 | 5.25 | 0.2 | 0.86 | 0 | 19.29 | 0 | x | x | Decreased |
| *21 | 0.1 | 0 | 0 | 0.5 | 0 | 0 | x | x | None |
| *22 | 0.06 | 0.3 | 0 | 0 | 0 | 0 | x | x | Unknown |
| *27 | 0.12 | 0 | 0.14 | 0 | 0.38 | 0 |  |  | Normal |
| *28 | 0.12 | 0.5 | 0.14 | 0 | 0 | 0 | x | x | Unknown |
| *29 | 2.64 | 0 | 0.29 | 0 | 9.83 | 0 | x | x | Decreased |
| *31 | 0.12 | 0.2 | 0.58 | 0 | 0 | 0 | x | x | None |
| *32 | 0.08 | 0.3 | 0.14 | 0 | 0 | 0 |  |  | Unknown |
| *33 | 0.18 | 0.6 | 0.29 | 0 | 0 | 0.1 | x | x | Normal |
| *34 | 0.02 | 0 | 0 | 0 | 0.08 | 0 |  |  | Normal |
| *35 | 1.48 | 4.67 | 2.59 | 0 | 0.23 | 0.61 | x | x | Normal |
| *36 | 0.26 | 0 | 0 | 0.2 | 0.83 | 0 |  |  | None |
| *39 | 0.08 | 0 | 0.14 | 0 | 0.08 | 0.2 |  | x | Normal |
| *40 | 0.24 | 0 | 0 | 0 | 0.91 | 0 | x | x | None |
| *41 | 6.07 | 8.75 | 5.91 | 3.77 | 1.59 | 11.86 | x | x | Decreased |
| *43 | 0.5 | 0.1 | 0 | 0 | 1.06 | 1.02 | x | x | Unknown |
| *45 | 0.88 | 0 | 0.29 | 0 | 3.18 | 0 | x | x | Normal |
| *46 | 0.16 | 0 | 0.14 | 0 | 0.53 | 0 | x | x | Normal |
| *49 | 0.1 | 0 | 0 | 0.5 | 0 | 0 | x |  | Decreased |
| *52 | 0.02 | 0 | 0 | 0.1 | 0 | 0 | x | x | Unknown |
| *56 | 0.02 | 0 | 0 | 0 | 0.08 | 0 | x | x | None |
| *59 | 0.06 | 0.2 | 0.14 | 0 | 0 | 0 | x | x | Decreased |
| *68 | 0.04 | 0 | 0 | 0 | 0.08 | 0.1 |  |  | None |
| *71 | 0.12 | 0 | 0 | 0.6 | 0 | 0 | x | x | Unknown |
| *82 | 0.06 | 0 | 0.43 | 0 | 0 | 0 | x | x | Unknown |
| *83 | 0.02 | 0 | 0 | 0 | 0 | 0.1 |  | x | Unknown |
| *84 | 0.02 | 0 | 0 | 0 | 0.08 | 0 |  |  | Unknown |
| *86 | 0.44 | 0 | 0 | 0 | 0 | 2.25 |  |  | Unknown |
| *99 | 0.04 | 0 | 0 | 0 | 0 | 0.2 | x | x | None |
| *106 | 0.32 | 0 | 0.14 | 0 | 1.13 | 0 | x | x | Unknown |
| *108 | 0.06 | 0.3 | 0 | 0 | 0 | 0 |  | x | Unknown |
| *111 | 0.16 | 0 | 0 | 0 | 0 | 0.82 | x | x | Unknown |
| *112 | 0.04 | 0 | 0 | 0 | 0 | 0.2 | x | x | Unknown |
| *113 | 0.16 | 0 | 0 | 0 | 0 | 0.82 | x | x | Unknown |
| *117 | 0.08 | 0.4 | 0 | 0 | 0 | 0 |  |  | Unknown |
| *121 | 0.02 | 0 | 0 | 0 | 0.08 | 0 |  |  | Unknown |
| *125 | 0.1 | 0 | 0 | 0 | 0.38 | 0 |  |  | Unknown |
| *139 | 0.02 | 0 | 0 | 0 | 0.08 | 0 |  |  | Unknown |
| *1x2 | 0.54 | 0.5 | 1.44 | 0.1 | 0.45 | 0.51 | x | x | Increased |
| *1x3 | 0.02 | 0 | 0 | 0 | 0.08 | 0 |  |  | Increased |
| *2x2 | 1.14 | 1.49 | 0.58 | 0.6 | 2.12 | 0.41 | x | x | Increased |
| *2x3 | 0.04 | 0.1 | 0 | 0 | 0.08 | 0 |  |  | Increased |
| *4x2 | 0.88 | 0.3 | 0.14 | 0 | 3.03 | 0 | x | x | None |
| *4x3 | 0.04 | 0 | 0 | 0 | 0.15 | 0 |  |  | None |
| *9x2 | 0.02 | 0.1 | 0 | 0 | 0 | 0 |  |  | Normal |
| *10x2 | 0.06 | 0 | 0 | 0.3 | 0 | 0 | x | x | Decreased |
| *17x2 | 0.02 | 0 | 0 | 0 | 0.08 | 0 |  | x | Normal |
| *29x2 | 0.1 | 0 | 0 | 0 | 0.38 | 0 |  |  | Normal |
| *35x2 | 0.02 | 0 | 0.14 | 0 | 0 | 0 |  |  | Increased |
| *43x2 | 0.04 | 0 | 0.14 | 0 | 0.08 | 0 |  |  | Unknown |
| *45x3 | 0.02 | 0 | 0 | 0 | 0.08 | 0 |  |  | Increased |
| *36+*10 | 7.15 | 0 | 0.14 | 34.13 | 0.08 | 1.23 | x | x | Decreased |
| *36+*36 | 0.04 | 0 | 0 | 0.2 | 0 | 0 |  |  | None |
| *68+*4 | 1.94 | 5.57 | 2.45 | 0 | 0.23 | 2.15 | x | x | None |
| *68+*68+*4 | 0.08 | 0.1 | 0.43 | 0 | 0 | 0 |  |  | None |
| *36+*36+*10 | 0.36 | 0 | 0 | 1.79 | 0 | 0 | x | x | Decreased |
| *36+*36+*36+*10 | 0.02 | 0 | 0 | 0.1 | 0 | 0 | x | x | Decreased |
| *13+*2 | 0.1 | 0.2 | 0.43 | 0 | 0 | 0 | x | x | Normal |
| *4.013+*4 | 0.14 | 0.7 | 0 | 0 | 0 | 0 |  | x | None |
| *1+*90 | 0.02 | 0 | 0 | 0.1 | 0 | 0 | x | x | Unknown |
| *36+*36+*83+*10 | 0.02 | 0 | 0 | 0.1 | 0 | 0 |  |  | Unknown |
| Unknown | 1.92 | 0.6 | 2.31 | 3.57 | 1.97 | 1.23 |  |  |  |
| % haplotypes overlapping the validation set | 96.1 | 97.4 | 96.5 | 95.9 | 95.1 | 96.1 |  |  |  |

We next assessed Mendelian consistency of the Cyrius calls in sequencing data from 597 trios (Table S2 and Supplementary Information). While the comparison above against truth genotypes allows for different haplotype phasing, the Mendelian consistency check is a more stringent check of the phasing of the star alleles when more than two copies of *CYP2D6* are present. Of the 572 trios with calls in all three family members, 561 (98.1%) are Mendelian consistent. All of the inconsistent trios could be resolved by changing the phasing - i.e. no proband had a called star allele that was not present in one of the parents. The majority (8/11) of the inconsistent cases are where the trio identified that two identical copies of *CYP2D6* should be on the same haplotype with the other haplotype having zero copy of *CYP2D6* (i.e. *Cyrius call *\*1/*1* vs. trio-based phasing *\*5/*1×2*). This Mendelian consistency check confirms the consistency of the genotypes across the pedigree but not the accuracy of the star alleles called. Combining the trio concordance tests with the accuracy tests performed above against truth genotypes provides confidence in the overall accuracy of the genotypes produced by Cyrius.

### *CYP2D6* haplotype frequencies across five ethnic populations

We next looked beyond the validation samples to study *CYP2D6* in the global population. For this, we analyzed the haplotype distribution by population (Europeans, Africans, East Asians, South Asians and admixed Americans) in 2504 unrelated 1kGP samples (Figure 5, Table 2, Table S3). Additionally, the predicted phenotype frequencies for these populations are illustrated in Figure S7. Cyrius made definitive diplotype calls in 2456 (98.1%) of the samples calling 52 distinct star alleles (The 48 no-calls are explained in Supplemental Information). Of these 52 star alleles, 40 overlapped star alleles that had been included in our validation data. These 40 alleles represent 96% of all the star alleles called in the 1kGP samples (Table 2).

**Figure 5.**
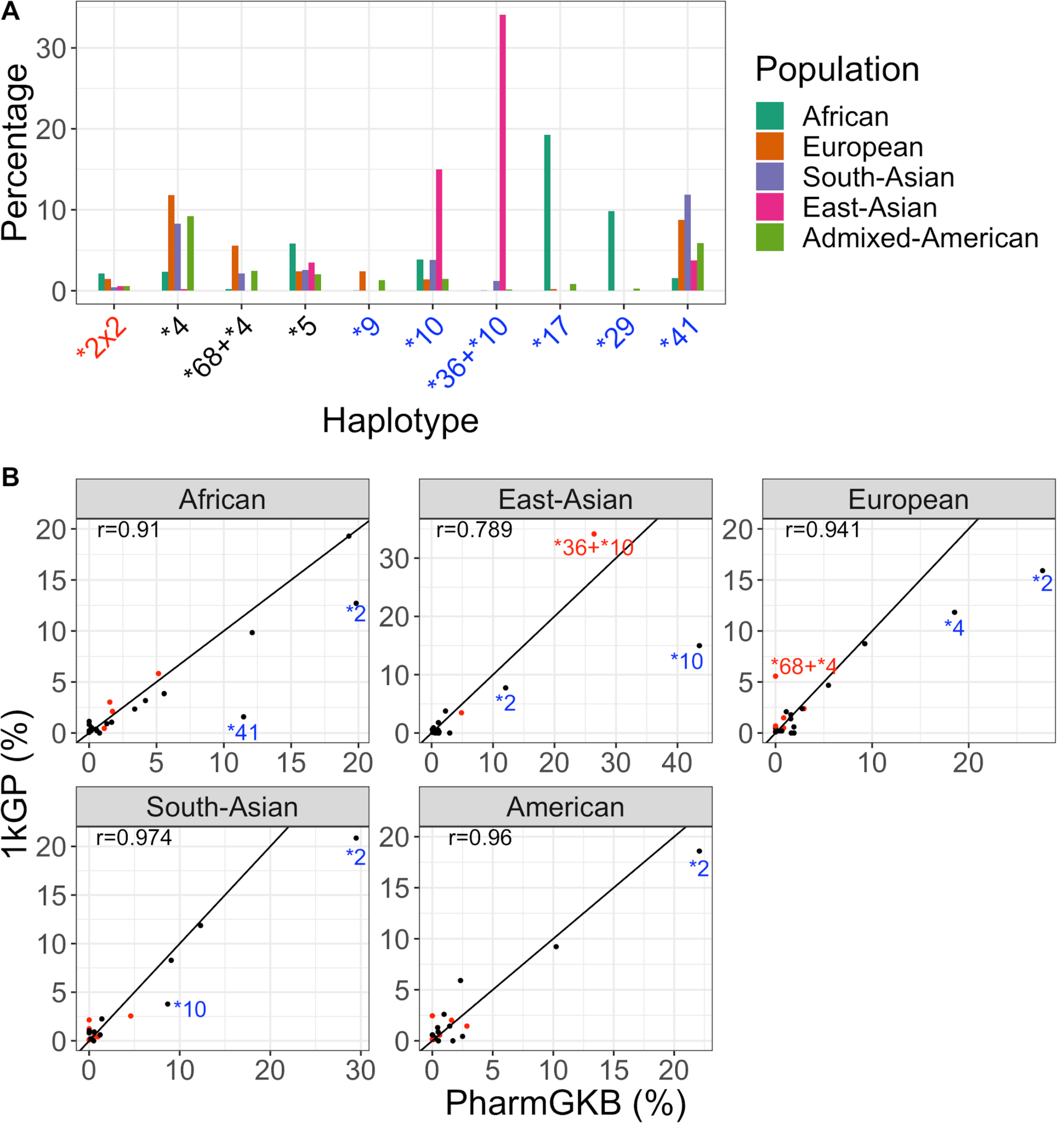
*CYP2D6* allele frequencies across five ethnic populations. **A**. Ten most common haplotypes with altered *CYP2D6* function. Those with increased function are labeled in red, those with no function in black and those with decreased function in blue. **B**. Comparison between 1kGP and PharmGKB frequencies. Each dot represents a haplotype with a frequency >=0.5% in either 1kGP or PharmGKB. SV-related haplotypes are marked in red, including the two haplotypes with the largest deviation (*\*36*+*\*10* in East-Asians and *\*68*+*\*4* in Europeans). Other haplotypes with deviated values are annotated in blue. A diagonal line is drawn for each panel. Correlation coefficients are listed for each population.

The haplotype frequencies mostly agree (correlation coefficient 0.79-0.97) with the summary of published allele frequencies in PharmGKB^6,31^ (Figure 5B, PharmGKB last accessed on 5/1/2020). While we report similar frequencies for *CYP2D6* deletion or duplication alleles as in PharmGKB, we report a higher frequency than PharmGKB for the SV-containing haplotype *\*36*+*\*10* in East-Asians and another SV *\*68*+*\*4* in Europeans (Figure 5B, red annotated dots). Previously reported frequencies of *\*36*+*\*10* in East-Asians fall into a wide range (10-35%)^32–37^, indicating the variability in CNV testing across assays. Additionally, *\*68* is often not interrogated in many studies, and it has been suggested that >20% of reported *\*4* alleles are actually in tandem with *\*68*^38,39^. Together, we estimate that the frequencies of haplotypes involving SVs are 38.6%, 11.2%, 11.4%, 6.8% and 7% in East-Asians, Europeans, Africans, Americans and South-Asians, respectively, and are 5.9%, 5.9%, 1.9%, 1.6% and 0.9% higher than reported in PharmGKB.

There are a few other star alleles for which we report a lower frequency than PharmGKB (Figure 5B, dots annotated in blue), highlighting the difficulty of merging data from multiple studies using different technologies^4^. These include *\*2* in all five populations. Since *\*2* is the default allele assignment for variant 2851C>T and 4181G>C unless additional variants defining other star alleles are interrogated, its frequency is likely overestimated in the literature^6^. Similarly, *\*10* is overestimated^4^ in East-Asians and South-Asians and *\*4* is overestimated^38^ in Europeans, particularly because a fraction of reported *\*10* or *\*4* alleles are *\*36*+*\*10* or *\*68*+*\*4*. Finally, we report a lower frequency for *\*41* in Africans, and *\*41* has not been consistently tested by its defining SNP across studies and is expected to be overestimated in Africans^31,40,41^.

## Discussion

We presented a new software tool, Cyrius, that can accurately genotype the highly complex *CYP2D6* region. Using 144 samples, including 8 with long read data, as an orthogonal validation dataset, we showed that Cyrius outperforms other *CYP2D6* callers, achieving 96.5% concordance versus 86.8% for Aldy and 84% for Stargazer. In particular, by using a novel CN calling approach, selecting a set of reliable *CYP2D6/CYP2D7* differentiating sites and accounting for possible mis-aligned reads, Cyrius is able to accurately identify star alleles with SVs, achieving 94.4% concordance compared to 87% for Aldy and 75.9% for Stargazer. Our comparison against the truth set allowed us to identify ways to improve the accuracy of Cyrius and after implementing those changes, we were able to increase the overall concordance to 99.3% (from 96.5%) and to 100% (from 94.4%) for the samples with SVs. We estimate that the star alleles miscalled in the validation data (*\*40, *46, *56* and *\*36* singleton) are only present in ∼0.68% of the population. Therefore, Cyrius’s accuracy is likely even higher in the population.

Across the 144 validation samples, we were able to confirm the accuracy of Cyrius across 40 different star alleles that represent roughly 96% of the star alleles in the pangenomic 1kGP population. In general, the allele frequencies we calculated for the five ethnic populations agree with previous studies for single copy star alleles. There are a number of limitations in the accuracy of the allele frequencies in PharmGKB because most studies test for a limited set of variants and there is often inadequate testing of CNVs^4,6^. WGS provides a promising option for building up more accurate population frequency databases because it assays all of the variants including CNVs and, combined with the right software, is able to resolve all of the known star alleles accurately. Furthermore, when new star alleles are added, it is easy to update allele frequencies by reanalyzing the same WGS data without having to design a new assay.

In our analysis of the 1kGP samples, Cyrius is able to call a definitive genotype in 98.1% of the samples. A future direction is to better understand the 1.9% of the samples that were not called and improve our algorithm so that it can also resolve these genotypes. For example, in samples where multiple haplotype configurations are possible, it could be useful to take a probabilistic approach to derive the most likely genotype given the observed variants. In addition, continuing to sequence and test more samples will help confirm our ability to genotype rare star alleles and will also identify additional variants that can be used to distinguish ambiguous diplotypes.

WGS provides a unique opportunity to profile all genetic variations for the entire genome but many clinically important regions/variants are beyond the ability of most secondary analysis pipelines. *CYP2D6* is among the difficult regions in the genome that are both clinically important and also require specialized informatics solutions to supplement generic WGS pipelines. Such targeted methods have already been applied successfully to some difficult regions, such as repeat expansions^28^ and the *SMN1* gene^20^ responsible for spinal muscular atrophy. The method employed in Cyrius can be applied to solve other paralogs that suffer from the same homology problem. We are currently extending this method to genotype other pharmacogenes with a paralog, *CYP2A6* and *CYP2B6*, and will apply this method to more genes in the future. With the continued development of more targeted methods like Cyrius, we can help accelerate pharmacogenomics and move one step closer towards personalized medicine.

## Availability of data and materials

Cyrius can be downloaded from: https://github.com/Illumina/Cyrius

The 1kGP data can be downloaded from https://www.ncbi.nlm.nih.gov/bioproject/PRJEB31736/ and https://www.ncbi.nlm.nih.gov/bioproject/PRJEB36890/.

WGS data for 70 GeT-RM samples can be downloaded from: https://www.ebi.ac.uk/ena/data/view/PRJEB19931.

For NA12878, NA24385, and NA24631, the PacBio Sequel II data is available in SRA under PRJNA540705, PRJNA529679, and PRJNA540706, and the Illumina data is available in ENA under PRJEB35491. For the remaining 5 samples with PacBio truth, the PacBio Sequel II data is available from http://ftp.1000genomes.ebi.ac.uk/vol1/ftp/data_collections/HGSVC2/working/.

## Acknowledgements

We thank the CDC Genetic Testing Reference Material Program (GeT-RM) for generating the consensus genotypes. We thank the New York Genome Center and the Coriell Institute for Medical Research for generating and releasing the 1kGP WGS data.

## Competing interests

XC, FS, NG, AM, CR, RJT, DRB and MAE are employees of Illumina Inc.

