## Supplementary Information for "Cyrius: accurate *CYP2D6* genotyping using whole genome sequencing data"

Xiao Chen[1], Fei Shen[1], Nina Gonzaludo[1], Alka Malhotra[1], Cande Rogert[1], Ryan J Taft[1], David R Bentley[2], Michael A Eberle[1]
1. Illumina Inc., 5200 Illumina Way, San Diego, CA, USA
2. Illumina Cambridge Ltd., Illumina Centre 19 Granta Park, Great Abington, Cambridge, UK

### Supplementary text

#### Star alleles targeted by Cyrius

Of all star alleles described by PharmVar (up to **139*), we removed two from our target list, **61* and **63* (both classified as "unknown function" and neither are included in GeT-RM). Both of these are *CYP2D6-CYP2D7* hybrids very similar to **36* with the fusion breakpoint slightly upstream. Importantly, these defined breakpoints occur in a part of the highly homologous Intron7-Exon8-Intron8 region between *CYP2D6/CYP2D7* (Figure S1) where the differentiating sites are not reliable (see below). Thus these two star alleles cannot be consistently distinguished from **36*, and they will be called as **36* by Cyrius.

#### Selecting reliable *CYP2D6/CYP2D7* differentiating sites

We identified 191 single nucleotide differences between *CYP2D6/CYP2D7* from the reference genome. Many of these base differences could be variable in the population, so not all of them could be used reliably to differentiate *CYP2D6* from *CYP2D7*. To identify reliable base differences, we focused on 1kGP samples where we called a CN(*CYP2D6*+*CYP2D7)* of 4 and thus we could assume that most likely these samples do not have structural variations, i.e. 2 copies of *CYP2D6* and 2 copies of *CYP2D7*. If a *CYP2D6/CYP2D7* base difference site is stable in the population, we expect that out of all reads that overlap this site in *CYP2D6* or *CYP2D7*, approximately half of the reads would have the *CYP2D6* base and we would call 2 copies of the *CYP2D6* base and 2 copies of the *CYP2D7* base in most samples. If the site is variable, i.e. *CYP2D6* base occurs in *CYP2D7* or vice versa, the CN of the *CYP2D6* base would often be called less than or greater than 2. Across the 191 sites, we queried the percentage of samples in which we called 2 copies of the *CYP2D6* base. Many sites showed a low percentage of samples with two copies of *CYP2D6* base (Figure S1) suggesting that the *CYP2D6/CYP2D7* base difference is not fixed in the population. Because we cannot be confident that *CYP2D6* does not have the *CYP2D7* base at these sites, they cannot be used to differentiate between the two genes. Relying on read alignments to these sites, as done by existing callers, would result in a large amount of noise when differentiating the two genes. We selected 117 highly stable sites with >98% of samples showing two copies of *CYP2D6* bases for *CYP2D6/CYP2D7* differentiation, which allowed us to gain the cleanest signal for calling SVs.

### Calling star alleles and assigning haplotypes

After calling SVs and small variants, Cyrius derives all possible star allele combinations that are consistent with the called variants. When there are multiple possible star allele combinations and one of them involves rare star alleles **34*, **39*, **4.009* and **139*, we favor the other more common combination(s). If there are more than one remaining star allele combination, Cyrius outputs all of them and flags the call as "More_than_one_possible_genotype".

To understand how often multiple star allele combinations could be consistent with the same set of variants, we examined the variants that would be present for all 10,731 pairwise combinations of the 146 star alleles (including some sub-alleles) interrogated by Cyrius. In this analysis, for 99.2% (10,650) of the pairwise star allele combinations, the variant sets were unique, 0.73% (78) were identical with one other star allele combination and 0.03% (3) were identical with two other star allele combinations. After removing combinations that result in the same combination of main star alleles and those that involve the 4 rare star alleles above, there are only 21 pairs (0.4%) of combinations that result in the same variant set. Therefore, the "More_than_one_possible_genotype" scenario only happens very rarely. In our population samples, we only observed one such combination, **1/*46* and **43/*45*, that is described below in "Discordant results with GeT-RM" and "No-calls made in 1kGP samples" sections.

After calling the individual *CYP2D6* copies, Cyrius assigns the star alleles into haplotypes. When there are more than two copies of *CYP2D6*, they are grouped into haplotypes based on known common tandem arrangements, e.g. **68+*4*, **36+*10, 36xN+*10, *13+*2 and *4.013+*4*.

If no common tandem arrangement is known, Cyrius reports all *CYP2D6* copies without assigning haplotypes, e.g. **1_*2_*68*, and flags the call as "Not_assigned_to_haplotypes".

### Aligners tested

The WGS aligners we have tested include BWA v0.7.15, Isaac v04.16.09.24 and DRAGEN v3.5.7b. For 2504 1kGP samples, we compared Cyrius calls for the same sample aligned with BWA and DRAGEN, and confirmed that the Cyrius genotype calls were identical.

### Discordant results with GeT-RM

Cyrius initially made five discordant calls (Table S1). Included amongst these discrepancies is the sample NA19908 (GeT-RM defined **1/*46*), where Cyrius called **1/*46* and **43/*45* as two possible diplotypes. Both of these two star allele combinations result in the same set of variants (77G>A, 1717G>A, 2851C>T and 4181G>C) and neither read phasing or population frequency analysis could rule out either genotype. The genotyping results from various assays that generated the GeT-RM consensus for this sample also showed disagreement between **1/*46* and **43/*45*, and the consensus was established by allele specific sequencing, highlighting the difficulty of these combinations (Table S4). Future sequencing of more samples of either diplotypes could help identify additional variants that distinguish the two.

In the remaining four samples where Cyrius calls were initially discordant with the truth, we were able to identify the causes and improve Cyrius to correctly call these star alleles. While we treat these four samples as miscalls for this study, we made improvements to Cyrius after seeing these samples, allowing us to call them accurately in the future.

First, in NA23275 (**1/*40*), the 18bp insertion defining **40* was initially missed as the insertion-containing reads were often not aligned properly and the inserted sequence together with the flanking sequence on one side were soft-clipped. Instead of just matching the alignment cigar, we modified Cyrius to look for this long insertion in the read sequences and improved the sensitivity for this insertion substantially.

Second, in HG03225 (**5/*56*), *CYP2D7*-derived reads were mis-aligned to *CYP2D6*, preventing the **56* defining variant from being called. We modified Cyrius to better distinguish and exclude mis-aligned reads in this region.

Third, in NA18565, we miscalled **10/*36x2* to be **10/*36+*10*, as did the other two callers. The depth profile shows a singleton form of **36* (*CYP2D6* with Exon 9 gene conversion, with REP6) in addition to the hybrid form of *36 (*CYP2D6-CYP2D7* hybrid, with REP7) (Figure S4). Cyrius was initially designed with the assumption that the base difference sites downstream of *CYP2D6* always share the same CN as the two base difference sites in Exon 9, so it used the consensus CNs of these sites to determine the CN of Exon 9. This case shows that when Exon 9 CN is smaller than the downstream sites, it suggests the presence of the singleton **36*. Another new variant type was learnt from the sample discussed in the next paragraph.

Lastly, in HG00421, we miscalled **10x2/*2* to be **2/*36+*10*, as did the other two callers. The depth profile shows a different SV, **10.003*, with the SV breakpoint located downstream of Exon 9 (between the base difference sites downstream of *CYP2D6* and the two base difference sites in Exon 9, Figure S4). This case shows that when Exon 9 CN is greater than the downstream sites, it suggests that the SV breakpoint is downstream of the gene, creating essentially a *CYP2D6* duplication instead of a hybrid (this is different from the most common breakpoints of *CYP2D6* duplications that fall in REP6/REP7). Cyrius was modified to include these two new variant types.

The last two samples (NA18565 and HG00421) have rare SV patterns so it is challenging to call such genotypes without seeing a known sample, as suggested by the fact that all three callers made the same wrong calls. As more samples with additional validated star alleles are sequenced, we can test the accuracy of Cyrius on these samples and improve the method as corner cases are identified.

### Mendelian consistency in 597 trios

We assessed Mendelian consistency of Cyrius calls in 597 1kGP trios (Table S2). Though for the accuracy comparison against the GeT-RM and PacBio validation data we did not require the phasing to be consistent with the 'truth' data, for this analysis we test both the star allele consistency and the phasing consistency. In 12 trios, the child is a no-call and in 13 trios, there is a no-call in one parent. The remaining 572 trios have definitive calls for all three samples. Out of these 572 trios, 561 (98.1%) are Mendelian consistent. Eight of the 11 inconsistent trios show an error in haplotype assignment, where, when there are two identical copies of *CYP2D6*, Cyrius defaults to assigning one copy to each haplotype and the true genotype has both copies on one haplotype, leaving a deletion on the other haplotype. For example: child HG03834 is called as **1x2/*4*, parent HG03833 is called as **1/*4* and parent HG03832 is called as **1/*1*. Based on the trio, parent HG03832 should be **1x2/*5*. The diplotype function is the same for **1x2/*5* and **1/*1*. In one other inconsistent trio (child HG00535), the discrepancy is between **10/*10+*36+*36* and **10+*36/*10+*36*, or **5/*10+*36* and **10/*36* (the diploype function is again the same). The remaining two inconsistent trios may reflect novel haplotypes or rare tandem arrangement (Table S2). Among the 13 trios where there is a no-call in one parent, the remaining duo is consistent except in one family: child HG02492 is called as **106/*2*, parent HG02491 is called as **1/*1* and parent HG02490 is a no-call. Based on this trio, we hypothesize that the child inherits **1* from one parent HG02491 and a novel haplotype from the other parent HG02490 that contains the merged variant set of **106* and **2* (3878G>A, 2851C>T and 4181G>C). This novel haplotype is also the cause of the no-call in parent HG02490.

### No-calls made in 1kGP samples

Out of 2504 1kGP samples, Cyrius did not call a definitive diplotype in 48 samples:

- 11 samples had an ambiguous depth pattern that resulted in failure to call CN(*CYP2D6*+*CYP2D7*) or failure to call SV.
- 4 samples have the same ambiguity between **1/*46* and **43/*45* as described in the GeT-RM sample NA19908 in “Discordant results with GeT-RM”. They are marked with the “More_than_one_possible_genotype” filter.
- 7 samples are flagged with the “Not_assigned_to_haplotypes” filter, indicating that they have more than two copies of *CYP2D6* (i.e. there is an SV) and Cyrius was not able to confidently phase them into haplotypes. For example, HG02778 and NA07347 are called as **1_*2_*68,* and we don’t have sufficient prior knowledge about which is the correct tandem arrangement (**68+*1* or **68+*2*, or **1+*2*).
- 26 samples had variant calls that did not match any of the known star alleles. For example, novel haplotypes can be easily inferred in samples where one haplotype is **5* and the other haplotype is a novel combination of variants (e.g. HG01551: c.4181G>C, c.2851C>T and c.77G>A; NA19314: c.4181G>C, c.2851C>T, c.983A>G, c.973C>A).

### Supplementary figures

#### Figure S1. *CYP2D6/CYP2D7* base difference sites show high variability in the population.

We identified 191 single nucleotide differences between *CYP2D6/CYP2D7* from the reference genome. We analyzed the CN call for the *CYP2D6* base at each of these 119 sites in 1kGP samples where we called a CN(*CYP2D6*+*CYP2D7* CN) of 4, i.e. most likely with 2 copies of *CYP2D6* and 2 copies of *CYP2D7*. The y axis shows the percentage of samples where we called the CN of the *CYP2D6* base as 2 across the 191 sites. The x axis shows genome coordinates in hg38. *CYP2D6* exons are drawn as orange boxes above the plot. We selected 117 highly stable sites with >98% (black horizontal line) of samples showing 2 copies of the *CYP2D6* base for *CYP2D6/CYP2D7* differentiation.

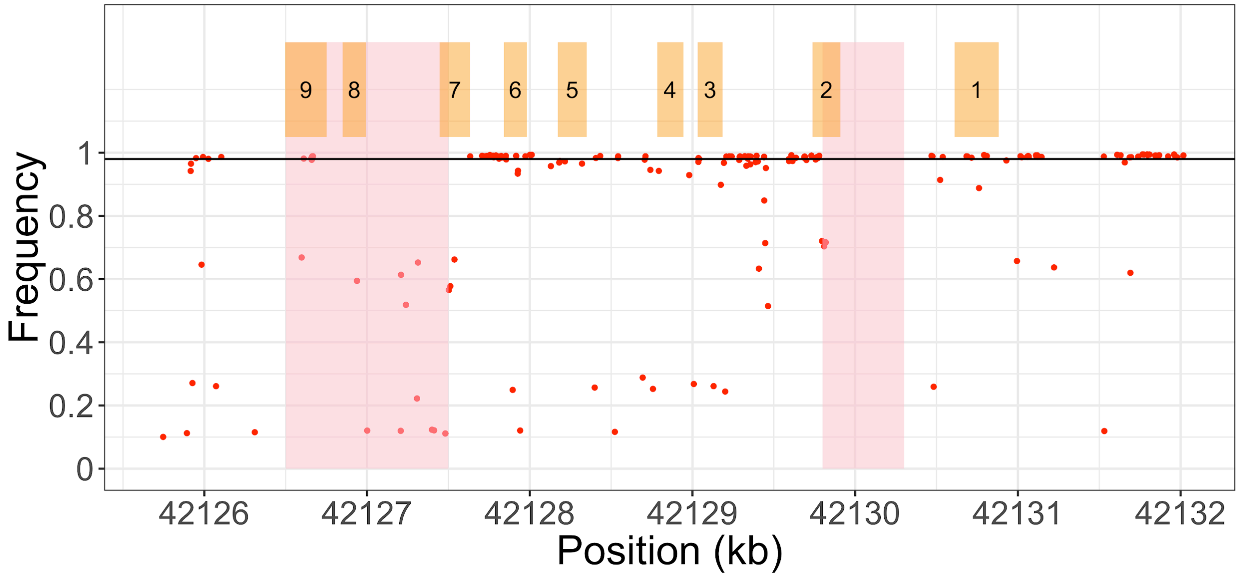

Figure S2. GeT-RM sample NA18519 (**106/*29*).

GeT-RM consensus is **1/*29* for NA18519, and the WGS-based genotype is **106/*29*. The IGV screenshot shows the variant-containing reads for variant g.3878G>C (g.42522916C>T, hg19) that defines **106*.

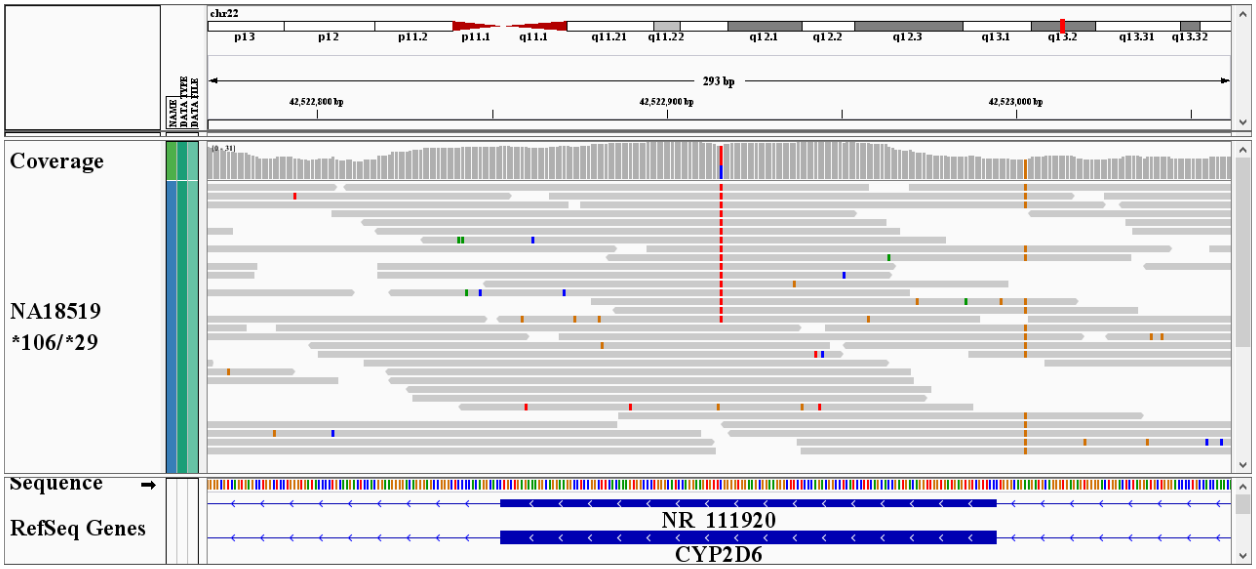

### Figure S3. Evidence of **68* in two GeT-RM samples (NA23874 and NA24008) where the three callers disagree with the GeT-RM consensus.

A. Depth plot as in Figure 2B. The Exon 1 region has a CN of 3, suggesting *CYP2D6*-*CYP2D7* hybrid. B. IGV screenshot of PacBio long read data showing reads that span the fusion breakpoint and thus align with clusters of mismatches. C. Realigning PacBio reads to sequence contigs representing the fusion, as in Figure 4, confirms the presence of the fusion in both samples.

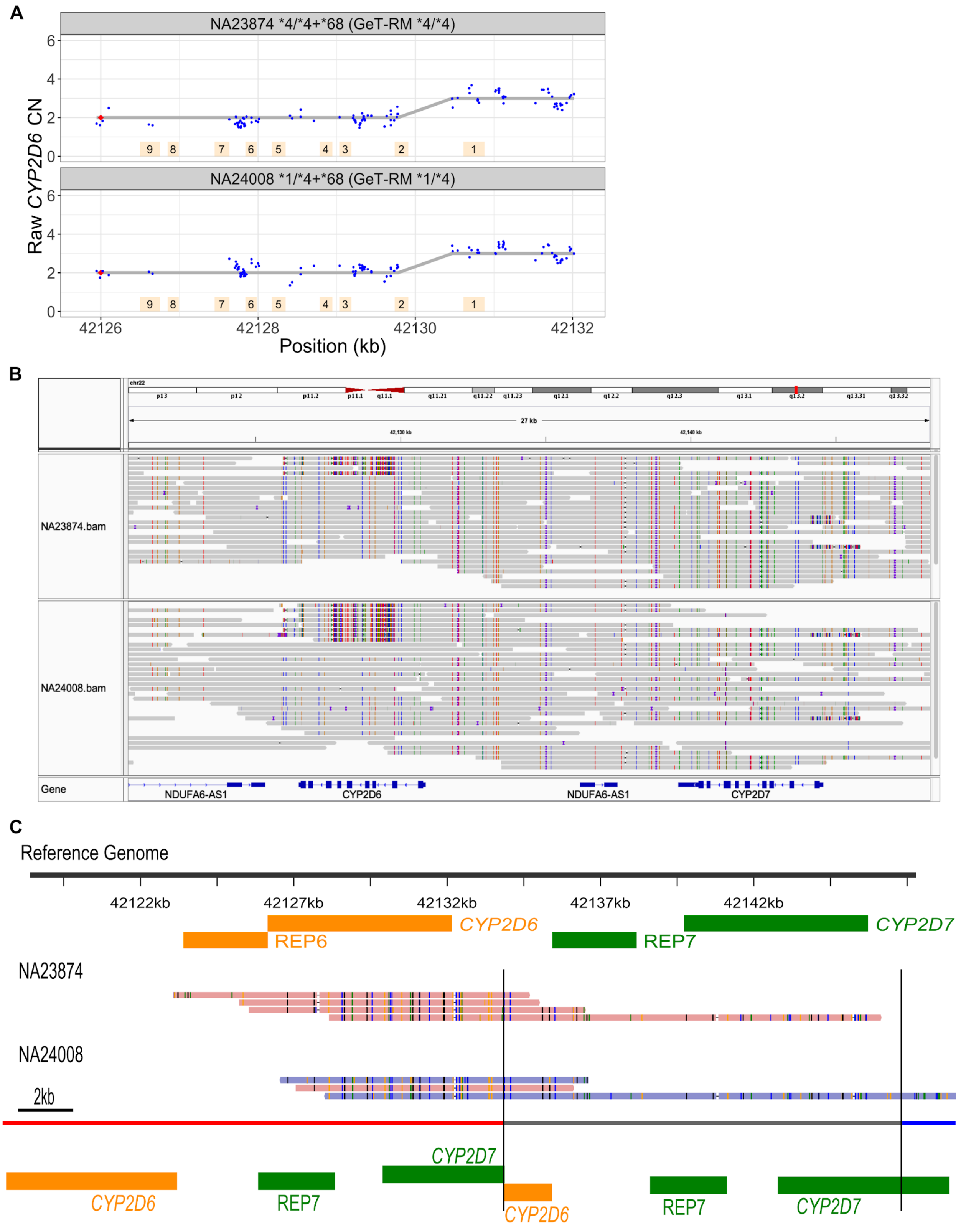

A
NA23874 *4/*4+*68 (GeT-RM *4/*4)
NA24008 *1/*4+*68 (GeT-RM *1/*4)
Raw CYP2D6 CN
0
2
4
6
9
8
7
6
5
4
3
2
1
42126
42128
42130
42132
Position (kb)
B
chr22
27 kb
42,130 kb
42,140 kb
NA23874.bam
NA24008.bam
Gene
NDUFA6-AS1
CYP2D6
NDUFA6-AS1
CYP2D7
C
Reference Genome
42122kb
42127kb
42132kb
42137kb
42142kb
CYP2D6
REP6
REP7
CYP2D7
NA23874
NA24008
2kb
CYP2D6
REP7
CYP2D7
CYP2D6
REP7
CYP2D7

### Figure S4. Depth plot for NA18565 and HG00421

Depth plot as in Figure 2B. NA18565 and HG00421 are both different from the commonly observed **36+*10* haplotype, as shown in NA18572 in the bottom panel, where the fusion breakpoint is upstream of the two base difference sites in Exon 9 so that the CN of Exon 9 is 2. For NA18565, where Cyrius initially called **10/*36+*10*, the CN of *CYP2D6* Exon 9 is 1 instead of 2, suggesting that there is a singleton form of **36* in addition to the hybrid form of **36.* Based on short read data, it is not clear whether the two copies of **36* are on the same haplotype. In HG00421, where Cyrius initially called **2/*36+*10*, the CN of *CYP2D6* Exon 9 is 3 instead of 2, suggesting that the duplication breakpoint is not within the gene but downstream of Exon 9 (between Exon9 and the base difference sites downstream of the gene), leaving the *CYP2D6* gene intact (**10.003*).

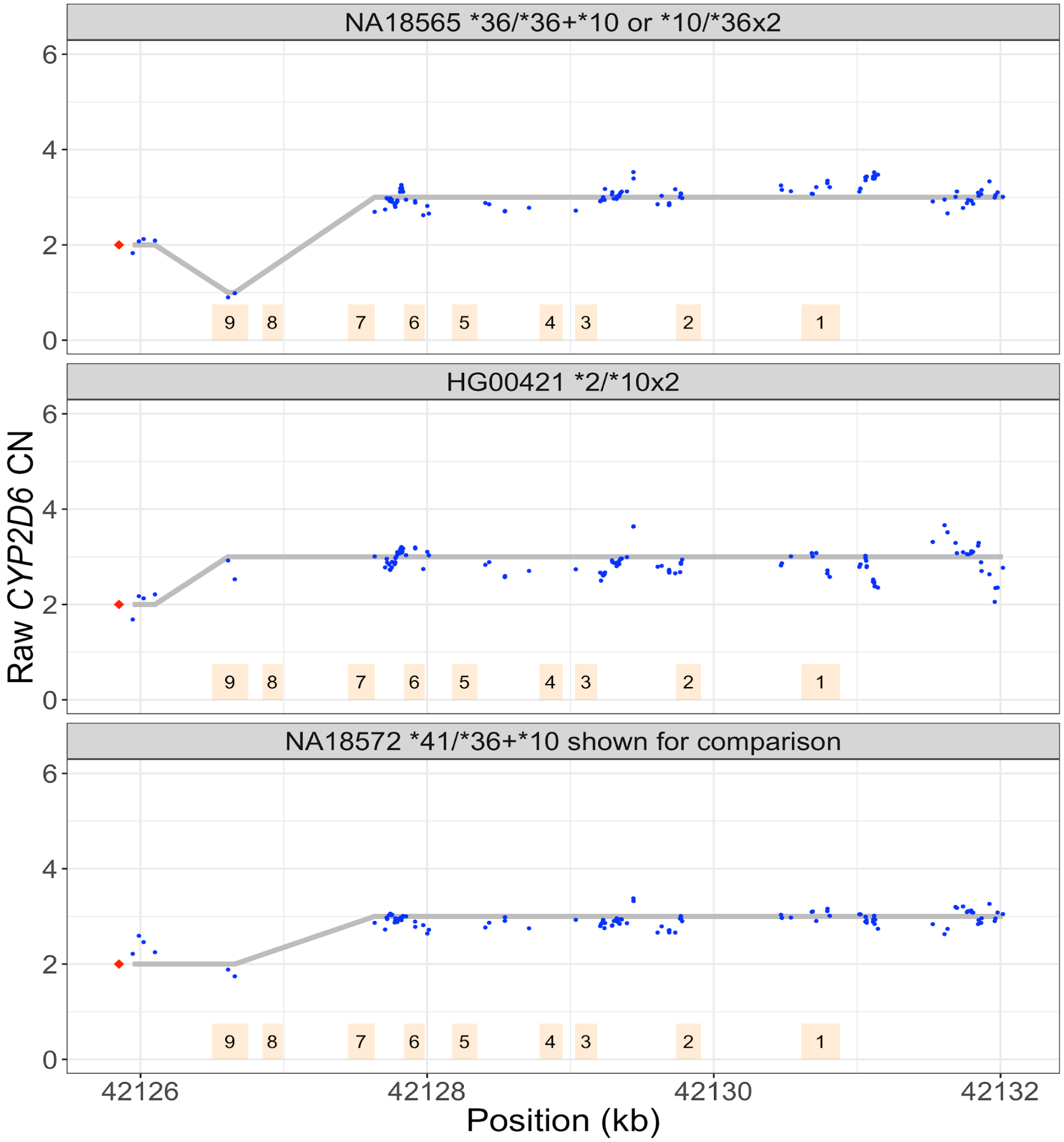

Figure S5. Depth plots for three samples with SVs where Cyrius calls remove uncertainty in GeT-RM consensus genotypes.

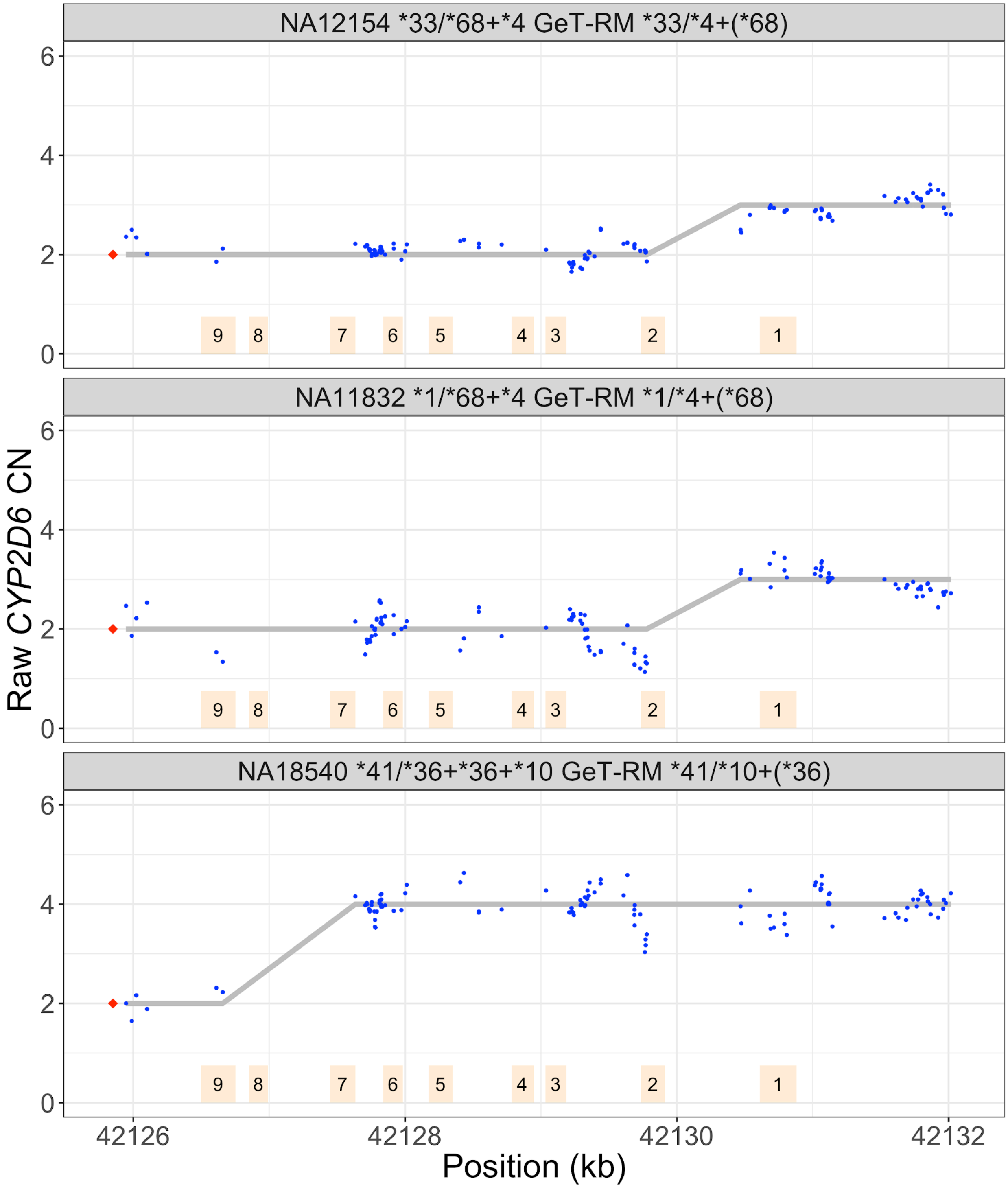

### Figure S6. Two samples without SVs where Cyrius calls remove uncertainty in GeT-RM consensus genotypes.

For NA07000, Cyrius calls **35/*9* and the GeT-RM consensus is **2 (*35)/*9*. For NA19143, Cyrius calls **45/*10* and the GeT-RM consensus is **2 (*45)/*10*. IGV screenshots show the variant-containing reads for variants that define **35* (g.31G>A, g.42526763C>T, hg19) and **45* (g.1717G>A, g.42525077C>T, hg19).

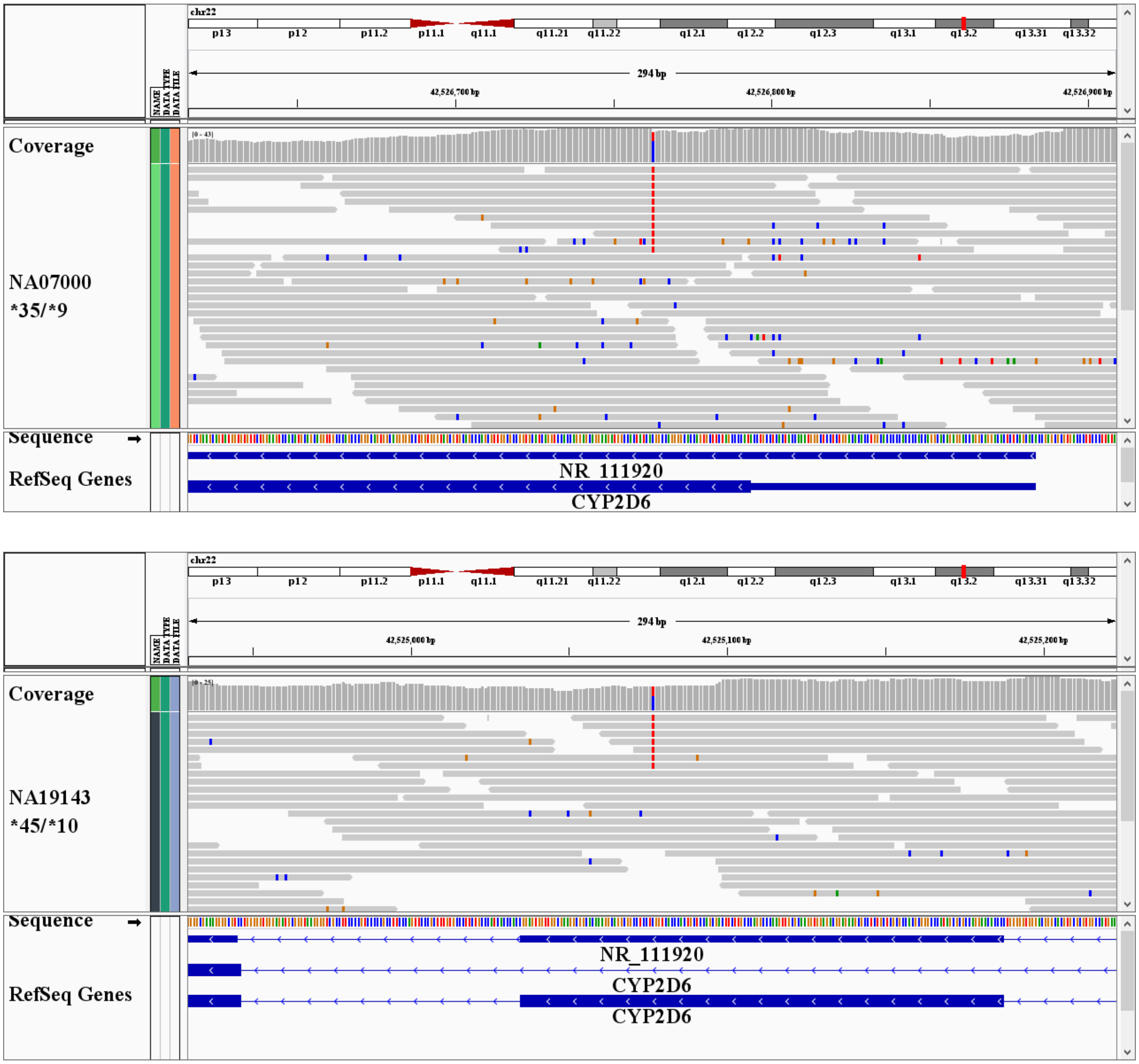

Figure S7. Distribution of predicted phenotypes across populations.

Following the latest CPIC guidelines, an activity score of 1 is predicted as an intermediate metabolizer.

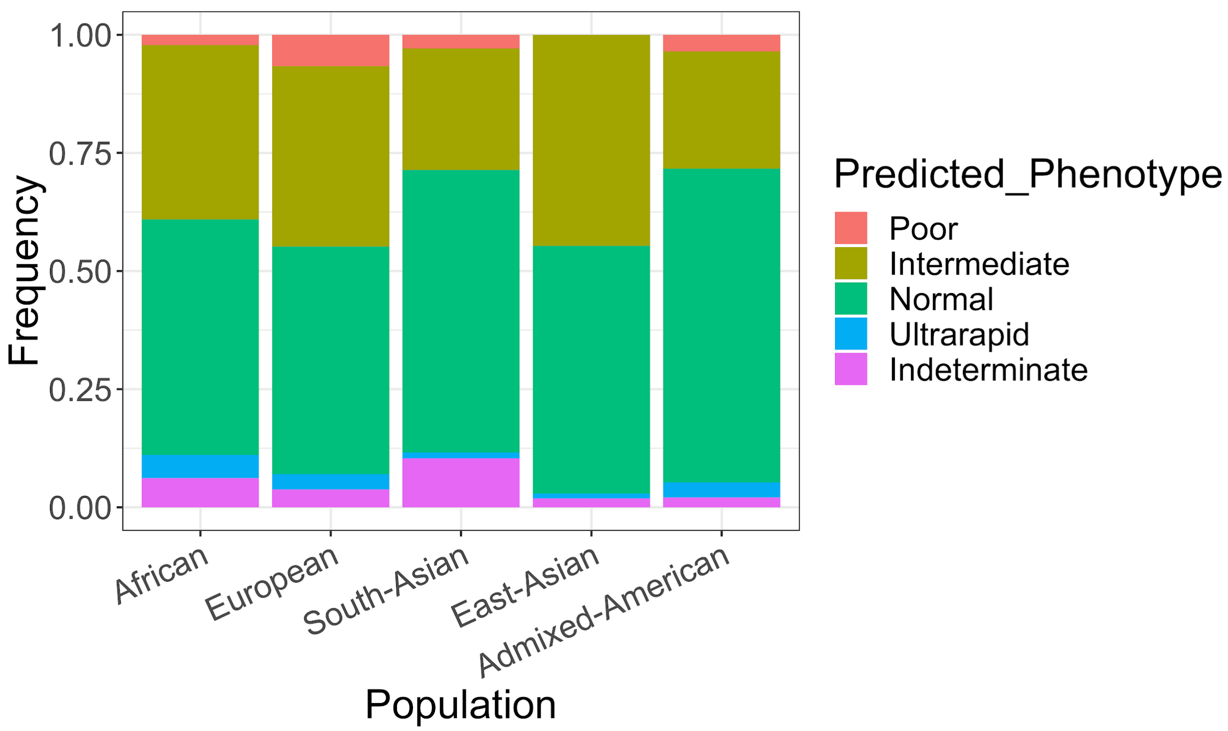

### Supplementary tables

Table S1. Cyrius/Aldy/Stargazer results against GeT-RM and PacBio truth. (Excel file)

Table S2. Cyrius calls in 597 trios. (Excel file)

Table S3. Cyrius calls in 1kGP samples. (Excel file)

Table S4. Additional information for GeT-RM samples where Cyrius calls were discrepant initially or helped remove GeT-RM uncertainty. (Excel file)
